# DishCam: an open-source 3D-printed darkfield imager for microbial colonies and small organisms on Petri dishes

**DOI:** 10.64898/2026.07.14.738485

**Authors:** David M. Versluis, Mattias Benninger, Devi Prasad Panigrahi, Anamul Hoque, Laura Machesky, Calvin Tiengwe, Robert Insall

## Abstract

Petri dishes are routinely photographed using handheld cameras under reflected light. While adequate for large, distinct colonies, this approach fails to capture individual organisms or colonies that are transparent, do not reflect light well, or have fine structures, and is unsuitable for high quality timelapses. Darkfield illumination provides superior contrast for organisms on agar surfaces, but existing solutions rely on improvised setups with inconsistent lighting, or on large and expensive benchtop instruments that cannot be housed inside incubators. We present DishCam, an open-source device consisting entirely of 3D-printed parts and inexpensive off-the-shelf electronics, controlled by a Raspberry Pi running custom open-source software. DishCam produces high-contrast darkfield images using angled LED illumination beneath a Petri dish. All components used are off-the-shelf or easily 3D-printed. The camera and lights can be fully controlled through Raspberry Pi using our software. We demonstrate the device across multiple biological systems: *Klebsiella aerogenes* colonies of differing densities, *Trypanosoma brucei* colonies doing social motility, *Dictyostelium discoideum* streaming aggregation, and *Caenorhabditis elegans* on feeding substrates. The total cost of the basic device is approximately £166 or $222, with the upgraded configuration costing approximately £413 or $552, which includes a high-quality telephoto lens, SSD storage and incubator-compatible cables.

## 1. Introduction

Darkfield illumination has long been used to visualise bacterial and eukaryotic colonies and other organisms on agar plates [Richards and Heijn, 1945, Wood, 1947]. These devices exploit the principle that light directed at a shallow angle around a transparent medium will refract off structures on or within that medium. A camera positioned above the medium can then register the scattered light but will not receive any unrefracted light from the light source. This renders the objects brightly visible against a dark background.

Many organisms can be studied more effectively using darkfield microscopy, while some are only visible under darkfield illumination. For example, Oberholzer et al. [2010], Imhof et al. [2014] and others used darkfield imaging to describe social motility in *Trypanosoma brucei* procyclic forms, demonstrating cooperative collective migrations that require sustained observation over days [DeMarco et al., 2020, Shaw et al., 2022]. Kuhn et al. [2024] subsequently quantified *T. brucei* social motility colony growth phases from such images. Similarly, Ingham and Ben-Jacob [2008] studied swarming and complex pattern formation in *Paenibacillus vortex* using real-time light microscopy. These studies highlight the need for consistent, automated imaging over extended periods.

Current approaches to Petri dish imaging suffer from several limitations. Handheld cameras under reflected light or improvised darkfield setups using angled lights (e.g. Oberholzer et al. [2010]) produce adequate single images of large, distinct colonies but fail to resolve fine structures. The images produced are also difficult to reproduce and have inconsistent illumination. Large benchtop imaging systems such as the Thermo Fisher iBright series (in e.g. Kuhn et al. [2024]) provide high contrast but are expensive, require fluorescent strains, and cannot be housed within incubators for live cell timelapse imaging. Microscope-based approaches offer high resolution but limited fields of view, and stitching multiple fields introduces artefacts. In recent years, open-source and 3D-printed imaging devices have begun to address the need for reproducible, low-cost colony imaging. Pi et al. [2022] introduced BAFFLE, a 3D-printable device for macroscopic quantification of fluorescent bacteria. While BAFFLE is well suited for multi-channel fluorescence imaging, it requires a relatively expensive DSLR camera and macro lens and is not designed for darkfield transmitted-light imaging of non-fluorescent colonies or organisms.

DishCam addresses these limitations by providing a compact, fully open-source darkfield imaging device that can be housed within standard laboratory incubators. The device is constructed entirely from 3D-printed parts and inexpensive off-the-shelf electronics, controlled by an unexpanded 1GB Raspberry Pi 5 with custom software that provides an optional touchscreen interface for live preview, single-frame capture, and automated timelapse acquisition.

## 2. Hardware description

DishCam consists of a 3D-printed frame supporting a Raspberry Pi High Quality Camera above a Petri dish, which is illuminated from below at a shallow angle using an LED strip, creating a darkfield effect (Fig. 1). The camera is controlled from a Raspberry Pi single-board computer, which can automate imaging via the included DishCam software.

**Figure 1:**
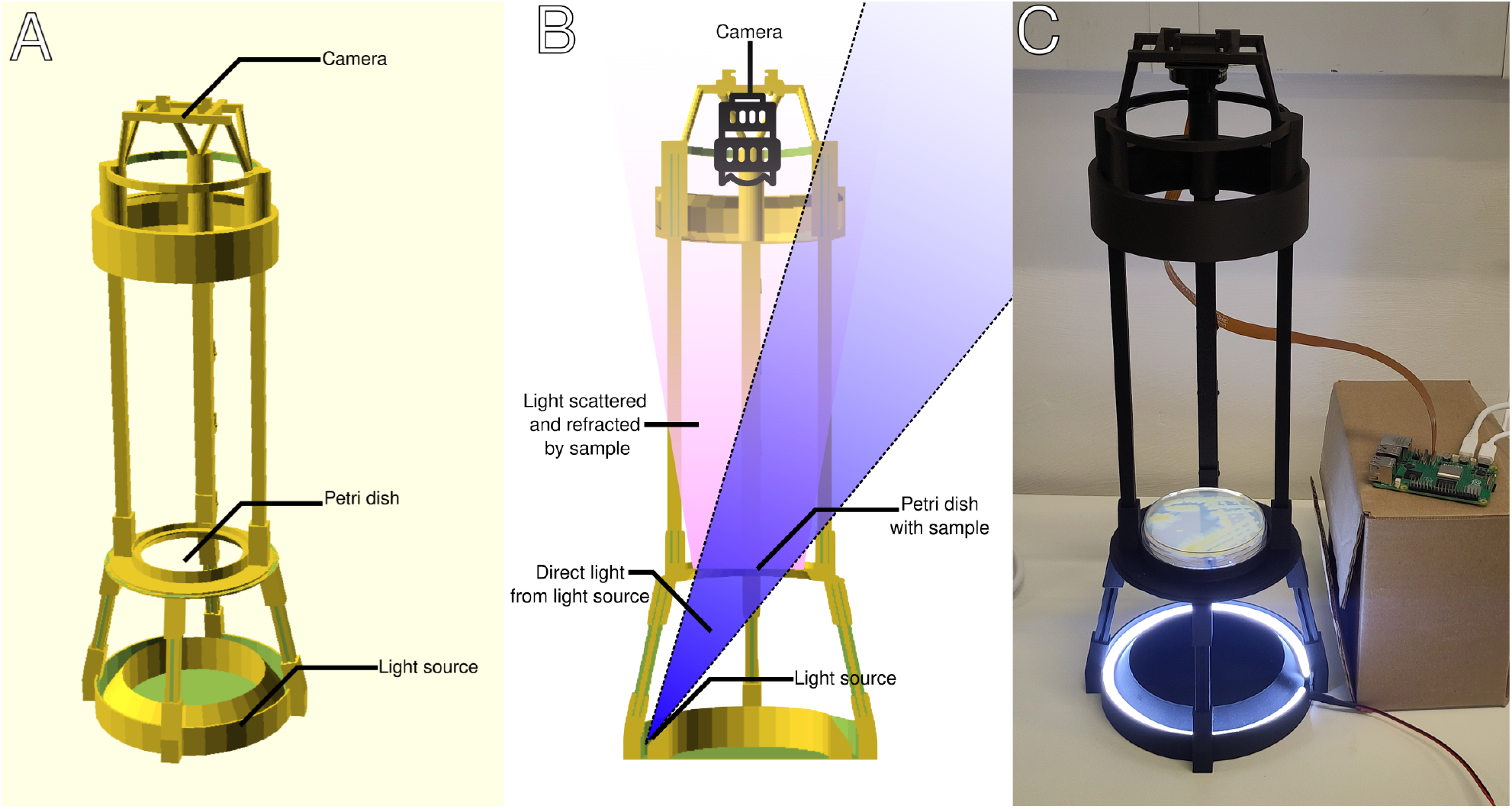
The DishCam basic configuration. (A) Annotated 3D render of the basic frame showing the three functional sections: camera attachment (top), Petri dish holder (middle), and light source holder (bottom). (B) 3D render cut in half, with light beams from the dish and one representative point of the LED strip visible, showing how light from the LEDs cannot directly be seen by the camera, but only when refracted by organisms on the dish. (C) Photograph of the assembled device with the Raspberry Pi Touch Display showing the DishCam software interface. The LED ring illuminates the Petri dish from below at an oblique angle while the camera captures images from above.

The basic device (Fig. 1) comprises three main functional elements. The upper part consists of a Raspberry Pi High Quality Camera mounted on a 3D-printed camera holder, supported above the Petri dish by three vertical beams. These beams are 300 mm long to ensure the field of view of the camera is flat from the centre to the edges of a Petri dish. The middle section holds a standard Petri dish (90 mm diameter) on a holder that centres the dish in the optical path. The lower section houses a 12 V LED strip on a conical mount, directing light upward at an oblique angle through the agar. This angle prevents direct light from reaching the camera, ensuring that only scattered light from structures on or in the agar contributes to the image. The angle of the lights, combined with the black 3D-printed background, creates excellent high-contrast darkfield images of microbial colonies, small organisms, or other small structures on the surface of the agar.

We have also created a more advanced version of the device (Fig. 2) that includes five independent optional additions, any of which can be incorporated into the basic design as needed. The first three additions can improve imaging quality; the other two additions make the device more portable and make imaging more user friendly.

**Figure 2:**
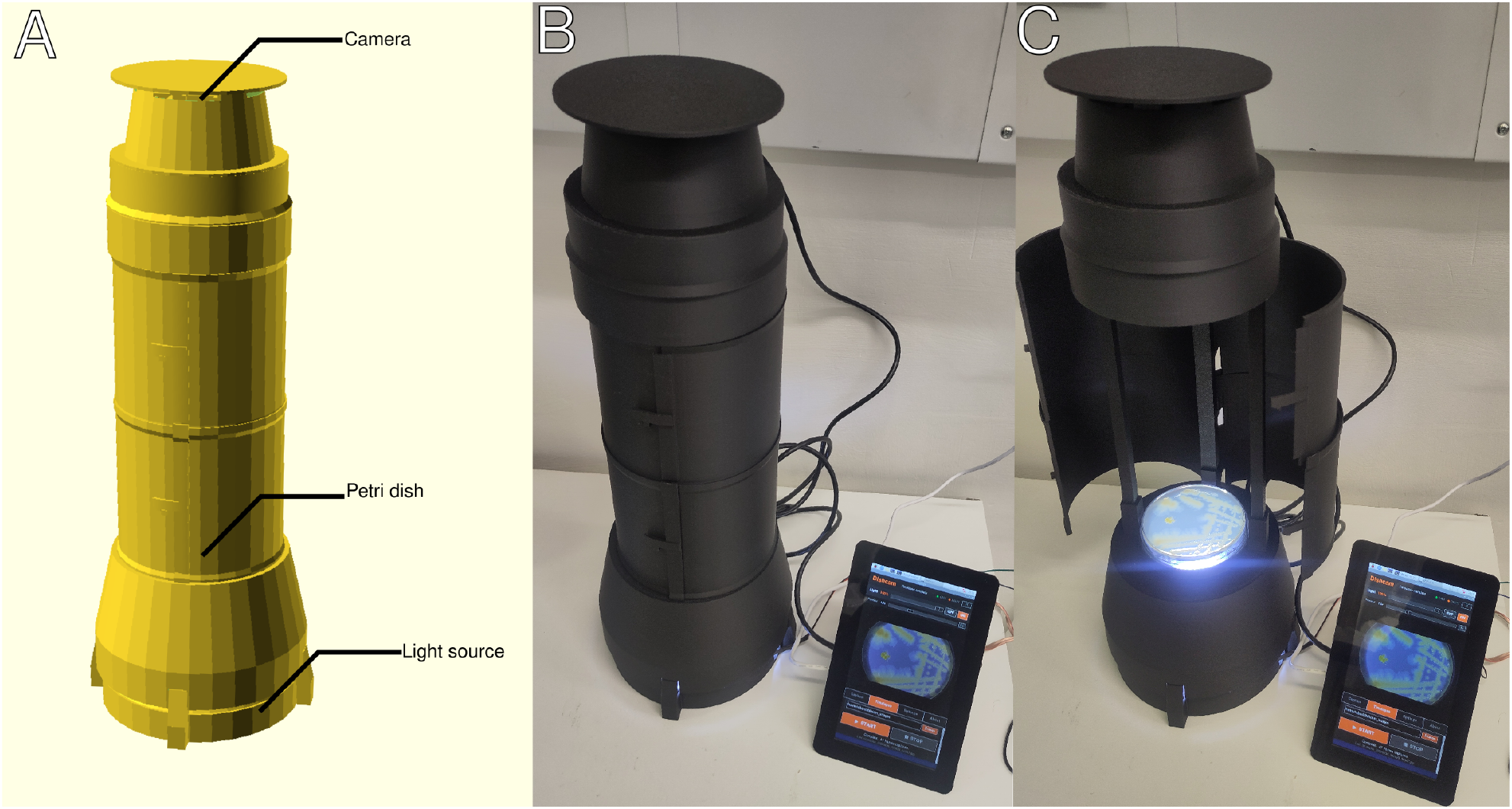
The DishCam advanced configuration with cylindrical light shades fitted. The hinged doors enclose the entire optical path from the LED ring to the camera, excluding ambient light. (A) Annotated 3D render (B) Photograph of the assembled device with closed doors. (C) Photograph of the assembled device with open doors.

1. A 16 mm telephoto Raspberry Pi High Quality Camera Lens can be used instead of the basic 12 mm lens. This allows for sharper images and also includes aperture control. To use the telephoto lens, the Raspberry Pi High Quality Camera with the default (C-CS) mount should be purchased instead of the version with the M12 mount.
2. The LED strip can be controlled by the Raspberry Pi through a MOSFET power controller, enabling dynamic modulation of the light intensity via hardware pulse-width modulation (PWM). This allows the lights to be dimmed to increase image quality and reduce heat and condensation during timelapse acquisition and permits the lights to be turned on only during image capture to further minimise thermal effects and save power.
3. A set of light shades can be fitted around the optical path from the light source to the camera, excluding environmental light and reflections to improve image quality and consistency.
4. The camera can be connected to the Raspberry Pi through an ethernet cable extension kit instead of the standard ribbon cable, allowing the imaging assembly to be placed at a greater distance from the Raspberry Pi. This is particularly useful in allowing the device to be placed inside an incubator while the Raspberry Pi remains outside.
5. A screen can be added (Raspberry Pi Touch Display 2). A printable frame provides support for both the MOSFET and the screen, and the dishcam software is set up to work well with the screen’s resolution and portrait orientation.

The included DishCam software (dishcam.py) provides a touchscreen-compatible graphical interface designed for the Raspberry Pi Touch Display 2 (720*×*1280 pixels) or any standard monitor. The software offers a live camera preview, single-image capture at full sensor resolution, and automated timelapse acquisition with configurable interval, total frame count, and per-frame light control. Light brightness is controlled via a slider with both software and hardware PWM output, and the interface includes camera exposure and gain settings with a preset optimised for fixed-light macro imaging.

## 3. Design files summary

| Design filename | File type | Open source license | Location of the file |
| --- | --- | --- | --- |
| Dishcam camera holder.stl | .stl file | CERN-OHL-P v2 | Available with the article |
| Dishcam dish holder.stl | .stl file | CERN-OHL-P v2 | Available with the article |
| Dishcam baseplate.stl | .stl file | CERN-OHL-P v2 | Available with the article |
| Dishcam bottom light shade.stl | .stl file | CERN-OHL-P v2 | Available with the article |
| Dishcam beam long doors.stl | .stl file | CERN-OHL-P v2 | Available with the article |
| Dishcam beam long normal.stl | .stl file | CERN-OHL-P v2 | Available with the article |
| Dishcam camera shade.stl | .stl file | CERN-OHL-P v2 | Available with the article |
| Dishcam door upper left.stl | .stl file | CERN-OHL-P v2 | Available with the article |
| Dishcam door upper right.stl | .stl file | CERN-OHL-P v2 | Available with the article |
| Dishcam door lower left.stl | .stl file | CERN-OHL-P v2 | Available with the article |
| Dishcam door lower right.stl | .stl file | CERN-OHL-P v2 | Available with the article |
| Pi screen support.stl | .stl file | CERN-OHL-P v2 | Available with the article |
| Dishcam frame.scad | CAD file (OpenSCAD) | CERN-OHL-P v2 | Available with the article |
| Pi screen support.scad | CAD file (OpenSCAD) | CERN-OHL-P v2 | Available with the article |
| dishcam.py | .py file | MIT License | Available with the article |

### Dishcam frame.scad

OpenSCAD source file for the 3D-printed components essential to the DishCam, including the base plate with LED strip mount, Petri dish holder with angled insert, camera holder with mounting plate and support beams (short prop beams and long camera beams). Also includes, as optional modules, the camera shade, and upper and lower light-shading doors with hinges and catches. The file is modular so that components can be modified or printed separately as desired. Requires a recent OpenSCAD nightly build, such as OpenSCAD-2026.06.21. All stls that start with “Dishcam” are from this .scad file

### Pi screen support.scad

OpenSCAD source file for the 3D-printed frame that supports the Raspberry Pi Touch Display 2 and MOSFET. The design is parametrised to allow different screen angles or sizes as desired. Requires threads.scad, available from https://github.com/rcolyer/threads-scad. “Pi screen support.stl” is from this file.

### dishcam.py

Python application providing a complete graphical user interface for camera control, live preview, single-frame capture, timelapse acquisition, and software and hardware PWM light control.

## 4. Bill of materials

A complete bill of materials is included as a separate file (Bill of Materials.csv). The basic configuration requires six components totalling approximately £166, $222, while the full advanced configuration adds up to ten optional components for a total of approximately £413, $553. All 3D-printed structural components are produced from PLA filament and are not included in the totals above.

## 5. Build instructions

### 5.1 3D Printing

All structural components can be 3D-printed from the single OpenSCAD source file (Dishcam frame.scad) or from the included .stl files. The openscad file also contains modules for each component, which can be rendered individually for printing. The 300 mm beams are the largest parts. They can be printed diagonally on build plates that are at least 240 mm per side. Alternatively, 300 mm MakerBeams can be purchased, which also provide superior strength. These do not allow for additional light shading to be attached. Optionally, a frame to hold the Raspberry Pi with screen can also be printed from the Pi screen support.stl file. This frame also provides an attachment point for the MOSFET light control.

We used a Bambu Lab P1S printer equipped with a 0.4 mm nozzle and a textured PEI print plate, and Bambu charcoal black PLA filament with the machine’s default settings. Any standard FDM printer with PLA would be sufficient to print these parts. For preparing gcode we used a standard 0.2 mm layer height throughout, with default tree supports enabled. Dark-coloured filament is recommended to minimise light reflections within the imaging chamber.

The components to be printed are: one base plate with LED strip mount, one Petri dish holder (mid-section), one camera holder with mounting plate, four short prop beams, one long camera beam with hinges for doors, and two long camera beams without hinges. For the advanced configuration with light shading, the lower and upper light shade doors (each printed as two halves) and the camera shade and light shade should also be printed. The beams can be replaced with metal MakerBeam beams (see bill of materials) for additional structural strength, or if the beams are too large to print on your printer. Metal beams do not allow the light shading doors to be attached.

### 5.2 Frame assembly

The 3D-printed components are assembled as follows:

1. Insert the LED strip into the bottom of the base plate. The LEDs should point upwards along the cone.
2. Insert the four short (100 mm) beams into the retainers on the base plate. The beams friction-fit. If the beams fit too tightly, they may be trimmed or sanded down with a small blade or sandpaper to achieve an appropriate fit.
3. Place the Petri dish holder (mid-section) onto the upper ends of the prop beams. Gently adjust the beams so they align smoothly with the Petri dish holder.
4. Screw the camera onto the camera mounting plate using the four mounting posts and attach with nuts (M2.5 screws and nuts). The camera holder includes a slot for the ethernet adapter if used.
5. Insert the three long (300 mm) camera beams into the beam connectors on the top of the mid-section.
6. Place the camera holder assembly onto the upper ends of the camera beams.

### 5.3 Light shading (optional)

The light shading consists of three kinds of parts: the bottom light shade, the doors, and the top light shade. The bottom light shade should be placed on the dish holder between steps 3 and 4 of assembly. The top light shade can simply be placed on top after the camera has been focused. The opening should be aligned with the camera cable. The door assemblies attach to the back camera beam via the printed hinges. Each door shade consists of two half-cylinders that open on hinges and close with a printed catch mechanism. The upper doors should be attached first, as far angled back as possible, and then moved to their forwards position. The bottom doors can then be attached in the same way. The doors can be opened for access to the Petri dish and closed during imaging to exclude ambient light.

### 5.4 Hardware setup

For the basic setup, simply connect the camera ribbon cable to the camera and Raspberry Pi, and the light cable to the 12V power adapter. The Raspberry Pi can be connected to peripherals and a monitor using the included cables.

When using an ethernet adapter to replace the ribbon cable, install according to the included installation instructions. If used, the Raspberry Pi SSD and Raspberry Pi Touch Display 2 should also be installed using the included instructions. A MOSFET can be used to have the light intensity be regulated by the Raspberry Pi. For the Gravity MOSFET we used (see bill of materials), the power adapters should be connected to VIN and GND on the MOSFET, and the lights connected to GND and VOUT. The three pins on the MOSFET cable should be connected to the Raspberry Pi as follows, top to bottom facing right: to pins 14 (GND), 17(3V3) and 12 (GPIO18) (Fig. 4).

If used, the Raspberry Pi can be attached to the back of the Raspberry Pi Touch Display 2. This can then be attached to a 3D printed frame (Fig. 3 B), along with the MOSFET, with the same M2.5 screws and bolts that can be used to attach the camera to the frame.

**Figure 3:**
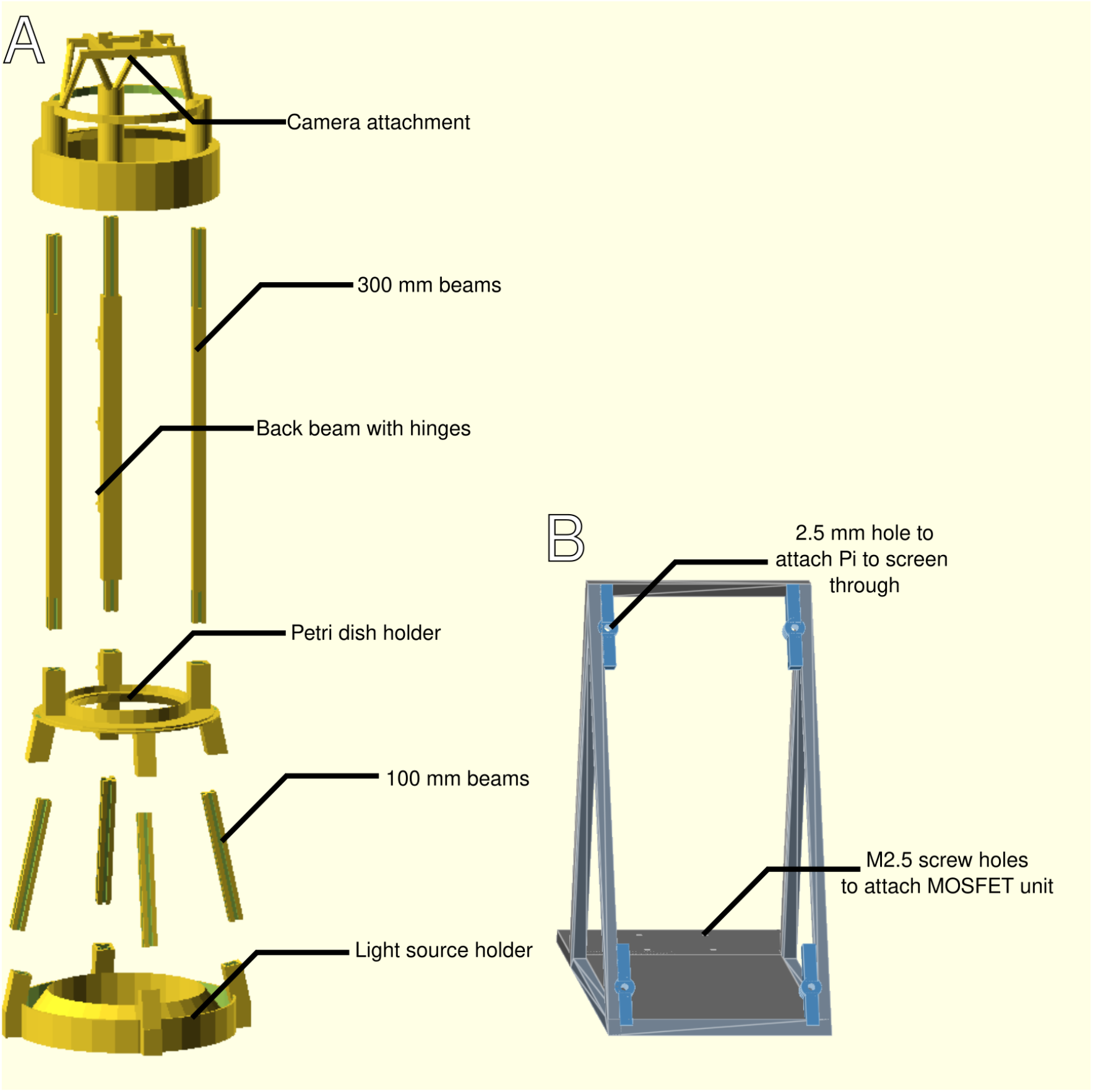
(A) Exploded view of the DishCam 3D-printed components, showing (from top to bottom): camera attachment, 300 mm beams, Petri dish holder, 100 mm beams, and light source holder. All components are printed from the single parametric OpenSCAD source file. (B) View of the optional 3D-printable frame to hold the screen and MOSFET unit. Not to scale with A.

**Figure 4:**
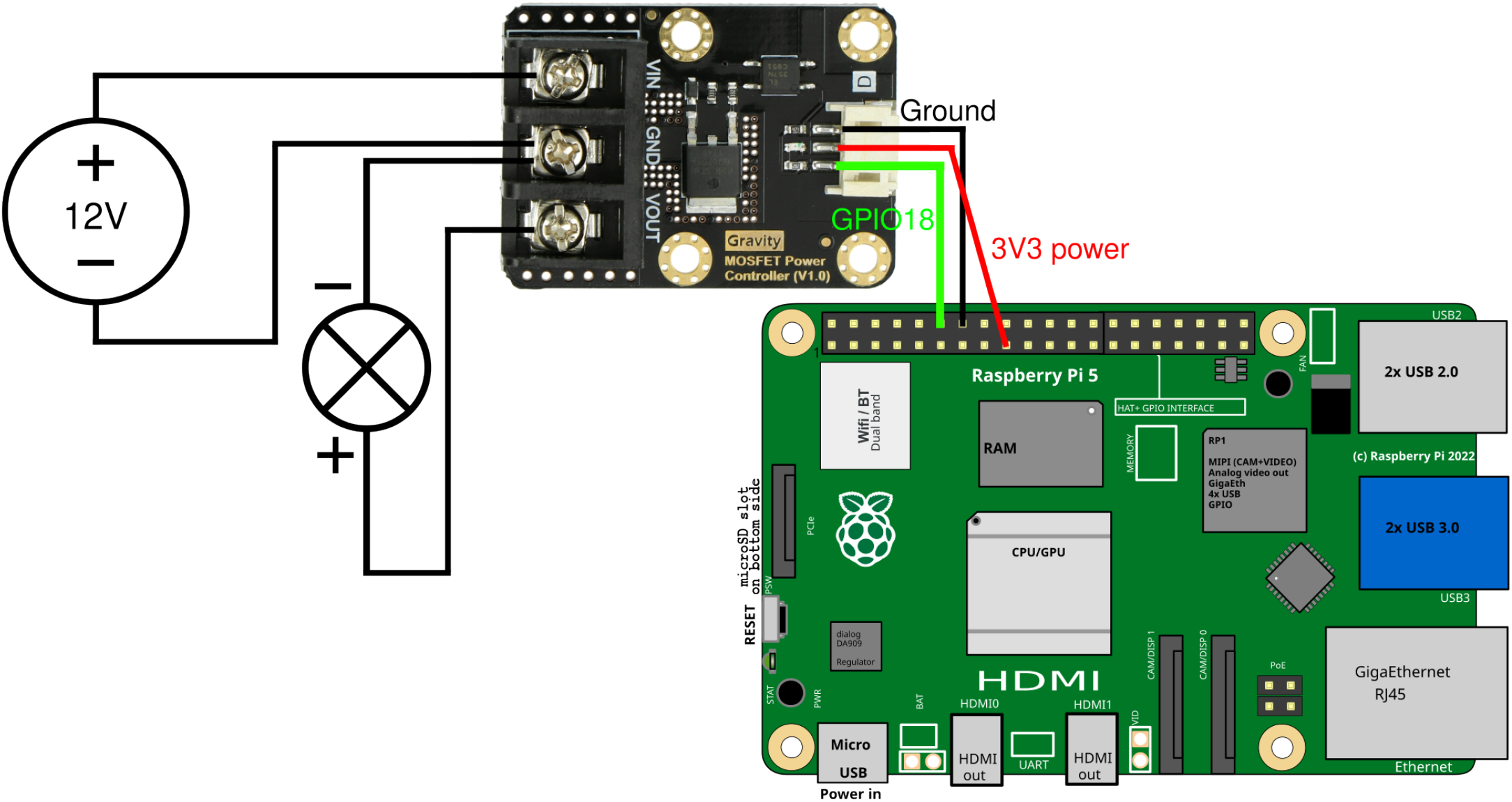
The connections to be made when using the MOSFET to have the Raspberry Pi regulate the lights.

### 5.5 Software setup

Install the required dependencies on the Raspberry Pi:

~~~
sudo apt install python3-picamera2 python3-pil python3-pil.imagetk
~~~

Copy dishcam.py to the Raspberry Pi. The software can then be launched with:

~~~
python3 dishcam.py
~~~

If the MOSFET is used to control the lighting, software PWM will be used by default. This may lead to flickering or uneven lighting depending on the processing load on the Raspberry Pi. To prevent this, hardware PWM may be enabled by adding the following line to `/boot/firmware/config.txt` on the Raspberry Pi:

~~~
dtoverlay=pwm,pin=18,func=2
~~~

The Raspberry Pi should then be rebooted and the software restarted to finalise setup.

## 6. Operating instructions

1. Place a Petri dish on the mid-section holder. The dish can be placed lid-down to reduce condensation on the imaging surface during timelapse acquisition.
2. Launch the DishCam software. A live preview will appear in the upper portion of the screen once the camera has initialised (indicated by the camera status dot turning green).
3. Turn the lights on by pressing the ON button on screen. Adjust the light brightness using the slider on the light bar to a level that allows for good contrast.
4. Focus the camera by rotating the ring on the lens. If using the optional 16 mm telephoto lens, the aperture (top ring) may also be adjusted. The live preview provides real-time feedback for focusing.
5. **Single capture:** Navigate to the Capture tab. Set the desired save folder and filename prefix, then press “CAPTURE PHOTO”. The image is saved as a full-resolution JPEG.
6. **Timelapse:** Navigate to the Timelapse tab. Set the interval between captures (hours, minutes, seconds), the total number of frames (0 for unlimited), and the filename prefix. Press “START” to begin acquisition. During timelapse, the software automatically turns the light on before each capture and off afterwards if a MOSFET is installed. Press “STOP” to end the timelapse at any time.
7. **Camera settings:** The Settings tab provides control over auto-exposure, auto-white-balance, manual exposure time, and analogue gain. A preset button loads settings optimised for our fixed-light timelapse imaging (AE off, AWB off, 1 s exposure, gain 1.0).

### Safety considerations

The device operates at 12V DC for the LED strip, 5V DC for the Raspberry Pi and 3V3 DC for the MOSFET, so no hazardous voltages are present. Some components may become warm during normal operation, but not dangerously so.

## 7. Results

### 7.1 Performance characteristics

The device captures a full 90 mm Petri dish in a single frame without stitching. The Raspberry Pi High Quality Camera provides a 4056*×*3040 resolution, and particularly with the 16 mm Raspberry Pi High Quality telephoto lens, very fine structures are resolvable. Timelapses are limited only by storage capacity (with the optional SSD, multi-week timelapses are feasible) and the minimum capture interval of approximately 2 seconds.

The device can fit inside an incubator. With the ethernet camera extension, only the camera and lights need to be placed inside the incubator. The principal limitations of the current design are that it provides darkfield transmitted-light imaging only, without support for fluorescence or reflected-light bright-field modes, and that the field of view is fixed to a single Petri dish. Image quality also depends on consistent agar thickness and surface flatness.

### 7.2 Case studies

We validated the DishCam across multiple biological systems to demonstrate its versatility and imaging quality. First, the device captured both fine-scale streaming structures and large-scale aggregation patterns of *D. discoideum* (Fig. 5 A, Video S1). The darkfield illumination provides sufficient contrast to resolve individual streaming arms at the macroscopic scale, without the need for fluorescent labelling.

**Figure 5:**
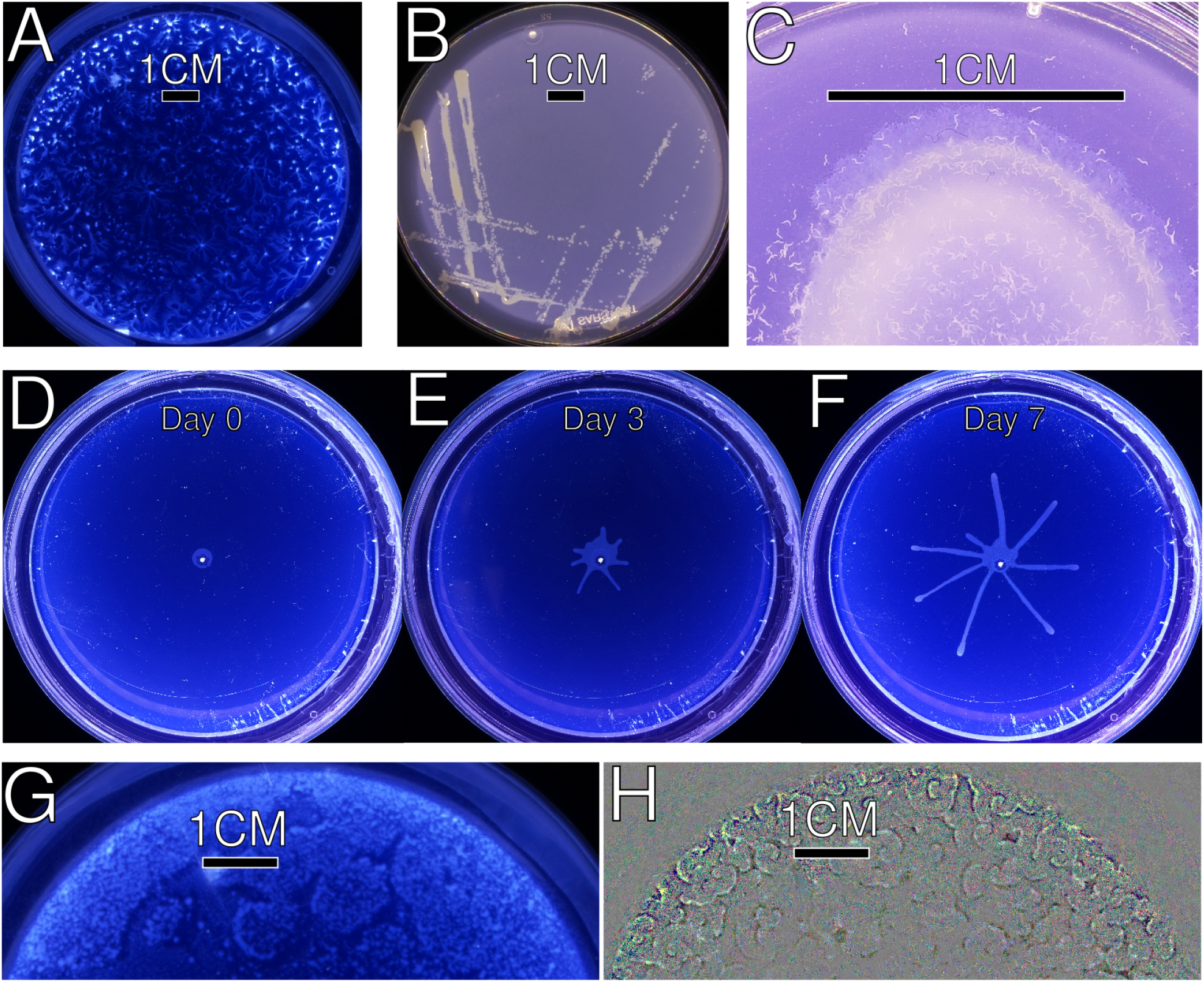
All images taken with DishCam. (A) *Dictyostelium* forming large streams on a 90mm Petri dish. (B) *Klebsiella aerogenes* spread on SM agar in a 90mm Petri dish at a gradually decreasing density. Individual colonies are clearly visible. (C) *C. elegans* on a bacterial feeding lawn. The darkfield illumination reveals both individual worms and the bacterial lawn structure. Close-up of a 30mm Petri dish. (D,E,F) Timelapse series of *T. brucei* social motility colony development on a 90mm Petri dish. (D) Shortly after inoculation, showing the initial colony at the centre of the plate. (E) Intermediate timepoint showing the emergence of radial colony projections. (F) Later timepoint with extended radial projections reaching toward the plate edge. All images are single full-resolution frames captured automatically during a 7 day timelapse. (G) *Dictyostelium* wave fronts in the early stage of aggregation on a 90mm Petri dish. (H) Analysis of differences between the frame in G and a frame one minute later, revealing finely structured spirals.

DishCam was subsequently used to image *Klebsiella aerogenes* on SM agar (Fig. 5 B). Even a thin semi-transparent bacterial layer can be seen clearly, as well as individual colonies.

We next used DishCam to image *C. elegans* on bacterial feeding lawns (Fig. 5 C, Video S2). Individual worms were clearly visible against the agar background, demonstrating the utility of the device for imaging organisms beyond microbial colonies. To utilise the device over longer periods, we made a timelapse of a procyclic *T. brucei* colony on agarose, replicating the social motility assay described by Oberholzer et al. [2010]. Timelapse imaging captured the full progression from initial inoculation through early colony projection to mature radial expansion (Fig. 5 D-F, Video S3), with fine temporal resolution revealing colony growth dynamics consistent with those described by Kuhn et al. [2024] (Figure S1), but without requiring fluorescent strains.

Finally, we returned to our images of *D. discoideum* and analysed them as they were pulsing during the early stage of aggregation [Brimson et al., 2025]. While the photos at this stage themselves do not clearly show the pulses (Fig. 5 G), the frequent high-resolution images allow us to compare subsequent images. This visualisation does clearly show the characteristic spiral pulsing waves (Fig. 5 H), which shows how the frequent high-resolution imaging allows for unexpected new insights.

## 8. Discussion

In conclusion, DishCam provides an accessible, low-cost darkfield imaging solution for Petri dishes, constructed entirely from 3D-printed parts and off-the-shelf electronics. We have demonstrated its versatility across multiple biological systems, including *Dictyostelium* streaming and aggregation, *C. elegans* on feeding lawns, and multi-day timelapse imaging of *T. brucei* social motility colonies. The device’s compact form factor allows it to be placed inside standard laboratory incubators, and its automated timelapse capability enables sustained observation that would be impractical with handheld or improvised setups. Furthermore, the high temporal resolution of the timelapse data can reveal phenomena not visible in individual frames. All hardware designs, 3D-printable files, and software are released as open source, enabling anyone to reproduce, modify, and extend the device for their own imaging needs.

## Supporting information

Bill of Materials, listing all required and optional components with pricing and sources

stl files of all components for 3d printing

Software (dishcam.py)

Scad files of both the imager and the optional screen support for rendering or further development

## 9. Supplemental Figure

**Supplemental figure S1.**
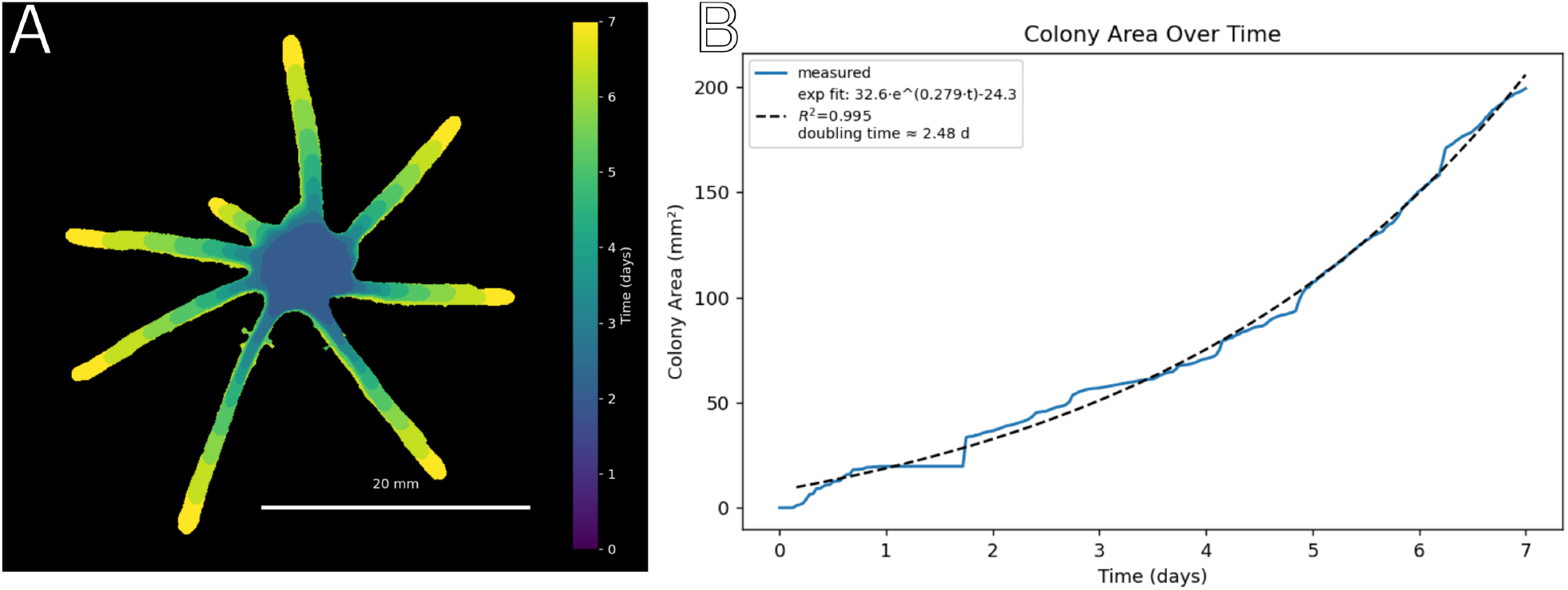
Visualisation and analysis of the *T. brucei* experiment from Fig. 5 and Video S3 using analysis methods first developed for fluorescent *T. brucei* colonies in Kuhn et al. [2024]. (A) Colony morphology per day (B) Colony size over time compared to fitted exponential growth curve

## 10. Supplemental Videos Video S1

*Dictyostelium* streaming and aggregating over 6 hours, imaged every minute.

**Video S2**

*C. elegans* on a feeding lawn moving around for 2 minutes, imaged every 2 seconds.

**Video S3**

*T. brucei* on SDM79 medium performing social motility at room temperature over 7 days, imaged every 2 minutes.

## 11. Acknowledgements

We thank Eric Lambie for providing the *C. elegans* plate. We are grateful to Sebastian Shaw and Isabel Roditi for advice on imaging *T. brucei*.

