## Supplementary material for "DishCam: an open-source 3D-printed darkfield imager for microbial colonies and small organisms on Petri dishes": Software (dishcam.py): dishcam.py

```
#!/usr/bin/env python3
"""
PetriCam – Raspberry Pi Petri Dish Imaging Application
Touchscreen GUI (720×1280) with live preview, single capture,
timelapse, and hardware-PWM light control on GPIO 18 (BCM).

Dependencies:
    sudo apt install python3-picamera2 python3-pil python3-pil.imagetk

Pin: Connect LED/MOSFET gate to GPIO 18 (BCM, physical pin 12).
"""

import logging
import os
import queue
import threading
import time
from datetime import datetime
from pathlib import Path

import tkinter as tk
from tkinter import filedialog, messagebox

# ──────────────────────────────────────────────────────────────────────────────
# Optional hardware imports – graceful fallback for off-Pi development
# ──────────────────────────────────────────────────────────────────────────────
try:
    from picamera2 import Picamera2
    CAMERA_AVAILABLE = True
except ImportError:
    CAMERA_AVAILABLE = False
    logging.warning("picamera2 not found – running in demo mode")

try:
    from PIL import Image, ImageTk, ImageEnhance
    PIL_AVAILABLE = True
except ImportError:
    PIL_AVAILABLE = False
    logging.warning("Pillow not found – preview disabled")

try:
    import RPi.GPIO as GPIO
    RPIGPIO_AVAILABLE = True
except ImportError:
    RPIGPIO_AVAILABLE = False


# ──────────────────────────────────────────────────────────────────────────────
# Constants
# ──────────────────────────────────────────────────────────────────────────────
APP_VERSION  = "1.0.0"
SCREEN_W     = 720
SCREEN_H     = 1180        # reduced from 1280 to leave room for the taskbar
PREVIEW_W    = 720
PREVIEW_H    = 430
PREVIEW_FPS  = 15          # frames per second in the live preview

GPIO_LIGHT   = 18          # BCM pin; hardware PWM channel 0 on all Pi models
PWM_FREQ_HZ  = 100         # default 100 Hz

DEFAULT_SAVE_DIR     = str(Path.home() / "dishcam_images")
DEFAULT_EXPOSURE_US  = 33_333    # ~30 ms (bright indoor / macro)
DEFAULT_GAIN         = 1.0

# Camera preset matching the original fixedlight.py settings
FIXEDLIGHT_PRESET = {
    "AeEnable":     False,
    "AwbEnable":    False,
    "ExposureTime": 1_000_000,   # 1 second
    "AnalogueGain": 1.0,
    "Contrast":     1.0,
    "Brightness":   0.0,
    "Saturation":   1.0,
    "Sharpness":    1.0,
}

# ──────────────────────────────────────────────────────────────────────────────
# Colour palette
# ──────────────────────────────────────────────────────────────────────────────
C = {
    "bg":       "#1A1A1A",   # main background
    "panel":    "#242424",   # header / tab bar
    "card":     "#2E2E2E",   # input cards
    "border":   "#3A3A3A",
    "orange":   "#FF6B00",   # primary accent
    "orhi":     "#FF8C33",   # orange hover
    "orlo":     "#CC5500",   # orange pressed / dim
    "white":    "#FFFFFF",
    "lgray":    "#CCCCCC",
    "mgray":    "#888888",
    "dgray":    "#444444",
    "red":      "#FF4444",
    "green":    "#44BB44",
    "amber":    "#FFAA00",
}

FF = "Helvetica"   # font family

logging.basicConfig(
    level=logging.INFO,
    format="%(asctime)s [%(levelname)s] %(name)s: %(message)s",
)
log = logging.getLogger("dishcam")


# ══════════════════════════════════════════════════════════════════════════════
# LightController
# ══════════════════════════════════════════════════════════════════════════════
class LightController:
    """
    Drives an LED / MOSFET on GPIO 18 via Linux sysfs hardware PWM.

    Uses only the sysfs interface — RPi.GPIO is deliberately avoided because
    GPIO.setup() resets the pin mux for GPIO 18, breaking hardware PWM and
    requiring a reboot to restore it.

    Requires dtoverlay=pwm,pin=18,func=2 in /boot/firmware/config.txt.
    Pi 5 uses pwmchip2/pwm2; Pi 4/3 use pwmchip0/pwm0. Auto-detected.

    All public methods are thread-safe.
    """

    # (chip, channel) pairs to try, in order: Pi 5 first, then Pi 4/3
    _PWM_CANDIDATES = [(0, 2), (0, 0), (1, 0)]

    def __init__(self, prefer_sysfs: bool = True):
        self._lock       = threading.Lock()
        self._brightness = 0.0    # 0.0 – 100.0 %
        self._enabled    = False
        self._gpio_pwm   = None
        self.backend     = "none"
        self._chip_path  = None
        self._pwm_path   = None
        self._freq_hz    = PWM_FREQ_HZ
        self._period_ns  = int(1e9 / PWM_FREQ_HZ)
        self.init_log    = []

        chips_found = [c for c, _ in self._PWM_CANDIDATES
                       if os.path.exists(f"/sys/class/pwm/pwmchip{c}")]
        self.init_log.append(
            f"sysfs chips present: {chips_found if chips_found else 'none'}")

        if not prefer_sysfs:
            self.init_log.append("sysfs skipped — using software PWM")

        for chip, channel in (self._PWM_CANDIDATES if prefer_sysfs else []):
            chip_path = f"/sys/class/pwm/pwmchip{chip}"
            if not os.path.exists(chip_path):
                continue
            pwm_path = f"{chip_path}/pwm{channel}"
            try:
                if not os.path.exists(pwm_path):
                    with open(f"{chip_path}/export", "w") as f:
                        f.write(str(channel))
                    time.sleep(0.1)
                else:
                    # Already exported (previous run) — disable before reconfiguring
                    try:
                        self._sysfs_write(f"{pwm_path}/duty_cycle", 0)
                        self._sysfs_write(f"{pwm_path}/enable", 0)
                    except Exception:
                        pass

                # period must be set before duty_cycle (default period=0 rejects writes)
                self._sysfs_write(f"{pwm_path}/period",     self._period_ns)
                self._sysfs_write(f"{pwm_path}/duty_cycle", 0)
                self._sysfs_write(f"{pwm_path}/enable",     1)
                self._chip_path = chip_path
                self._pwm_path  = pwm_path
                self.backend    = f"sysfs-pwmchip{chip}/pwm{channel}"
                msg = f"HW PWM ok: pwmchip{chip}/pwm{channel} @ {self._freq_hz} Hz"
                log.info("Light: %s", msg)
                self.init_log.append(msg)
                break
            except Exception as exc:
                msg = f"pwmchip{chip}/pwm{channel} failed: {exc}"
                log.warning("Light: %s", msg)
                self.init_log.append(msg)

        # ── RPi.GPIO software PWM fallback ────────────────────────────────────
        if self.backend == "none" and RPIGPIO_AVAILABLE:
            try:
                GPIO.setmode(GPIO.BCM)
                GPIO.setup(GPIO_LIGHT, GPIO.OUT)
                self._gpio_pwm = GPIO.PWM(GPIO_LIGHT, self._freq_hz)
                self._gpio_pwm.start(0)
                self.backend = "rpigpio"
                msg = f"SW PWM (RPi.GPIO) on GPIO {GPIO_LIGHT} @ {self._freq_hz} Hz"
                log.info("Light: %s", msg)
                self.init_log.append(msg)
            except Exception as exc:
                msg = f"RPi.GPIO PWM failed: {exc}"
                log.warning("Light: %s", msg)
                self.init_log.append(msg)

        if self.backend == "none":
            msg = "No PWM backend available (demo mode)."
            log.warning("Light: %s", msg)
            self.init_log.append(msg)

    # ── public API ────────────────────────────────────────────────────────────

    def set_brightness(self, pct: float) -> None:
        with self._lock:
            self._brightness = max(0.0, min(100.0, float(pct)))
            self._apply()

    def turn_on(self) -> None:
        with self._lock:
            self._enabled = True
            self._apply()

    def turn_off(self) -> None:
        with self._lock:
            self._enabled = False
            self._apply_duty(0.0)

    def get_brightness(self) -> float:
        return self._brightness

    def is_on(self) -> bool:
        return self._enabled

    def set_frequency(self, hz: int) -> bool:
        hz = max(10, min(100_000, int(hz)))
        with self._lock:
            new_period_ns = int(1e9 / hz)
            try:
                if self._pwm_path:
                    self._sysfs_write(f"{self._pwm_path}/duty_cycle", 0)
                    self._sysfs_write(f"{self._pwm_path}/enable",     0)
                    self._sysfs_write(f"{self._pwm_path}/period",     new_period_ns)
                    self._freq_hz   = hz
                    self._period_ns = new_period_ns
                    self._sysfs_write(f"{self._pwm_path}/enable", 1)
                    self._apply()
                    log.info("Light: frequency changed to %d Hz", hz)
                    return True
                elif self._gpio_pwm:
                    self._gpio_pwm.ChangeFrequency(hz)
                    self._freq_hz   = hz
                    self._period_ns = new_period_ns
                    log.info("Light: frequency changed to %d Hz", hz)
                    return True
            except Exception as exc:
                log.warning("Light: set_frequency error: %s", exc)
        return False

    def cleanup(self) -> None:
        with self._lock:
            try:
                if self._pwm_path:
                    # Disable but leave channel exported so re-init works without reboot
                    self._sysfs_write(f"{self._pwm_path}/duty_cycle", 0)
                    self._sysfs_write(f"{self._pwm_path}/enable", 0)
                elif self._gpio_pwm:
                    self._gpio_pwm.stop()
                    # Note: GPIO.cleanup() deliberately NOT called — it resets the
                    # pin mux for GPIO 18, breaking sysfs HW PWM until reboot.
            except Exception as exc:
                log.warning("Light cleanup error: %s", exc)

    # ── private ───────────────────────────────────────────────────────────────

    def _apply(self) -> None:
        self._apply_duty(self._brightness if self._enabled else 0.0)

    def _apply_duty(self, duty: float) -> None:
        try:
            if self._pwm_path:
                self._sysfs_write(f"{self._pwm_path}/duty_cycle",
                                  int(self._period_ns * duty / 100.0))
            elif self._gpio_pwm:
                self._gpio_pwm.ChangeDutyCycle(duty)
        except Exception as exc:
            log.warning("Light duty set error: %s", exc)

    @staticmethod
    def _sysfs_write(path: str, value) -> None:
        with open(path, "w") as f:
            f.write(str(value))


# ══════════════════════════════════════════════════════════════════════════════
# CameraController
# ══════════════════════════════════════════════════════════════════════════════
class CameraController:
    """
    Wraps picamera2.  Preview runs as RGB888 at PREVIEW_W×PREVIEW_H.
    Still captures switch to full-sensor resolution then return to preview.
    Timelapse captures stay in preview mode (no mode switch) for reliability.
    """

    def __init__(self):
        self._cam         = None
        self._lock        = threading.Lock()
        self._running     = False
        self.available    = False
        self._preview_cfg = None
        self._still_cfg   = None
        # Picamera2() is NOT called here – call start() from a background thread.

    def start(self) -> None:
        """
        Initialise and start the camera.  Call this from a background thread
        so the GUI is never blocked, even if libcamera takes several seconds.
        """
        if not CAMERA_AVAILABLE:
            log.warning("Camera: picamera2 not available (demo mode)")
            return
        try:
            self._cam = Picamera2()

            self._preview_cfg = self._cam.create_preview_configuration(
                main={"size": (PREVIEW_W, PREVIEW_H), "format": "BGR888"},
            )
            self._still_cfg = self._cam.create_still_configuration()

            self._cam.configure(self._preview_cfg)
            self._cam.start()
            self._running = True
            self.available = True
            time.sleep(2)   # allow AE/AWB to converge
            log.info("Camera: started (sensor %s)", self._cam.sensor_resolution)
        except Exception as exc:
            log.error("Camera start failed: %s", exc)

    def capture_frame(self):
        """Return the latest preview frame as a numpy array (H×W×3 RGB), or None."""
        if not self._running:
            return None
        try:
            return self._cam.capture_array("main")
        except Exception as exc:
            log.debug("Preview frame error: %s", exc)
            return None

    def capture_still(self, filepath: str, on_done) -> None:
        """
        Capture full-resolution still in a worker thread.
        on_done(success: bool, filepath: str) is called from the worker thread;
        caller must use root.after() if updating tkinter.
        """
        def _worker():
            ok = False
            try:
                with self._lock:
                    self._cam.switch_mode_and_capture_file(
                        self._still_cfg, filepath, wait=15
                    )
                ok = True
                log.info("Captured still: %s", filepath)
            except Exception as exc:
                log.error("Still capture failed: %s", exc)
            finally:
                on_done(ok, filepath)

        threading.Thread(target=_worker, daemon=True).start()

    def capture_timelapse_frame(self, filepath: str) -> bool:
        """
        Capture a JPEG in the current preview mode (no mode switch).
        Safe to call from a background thread.
        """
        if not self._running:
            return False
        try:
            with self._lock:
                self._cam.switch_mode_and_capture_file(
                        self._still_cfg, filepath, wait=15
                    )
            log.debug("Timelapse frame: %s", filepath)
            return True
        except Exception as exc:
            log.error("Timelapse capture error: %s", exc)
            return False

    def apply_controls(self, ctrl: dict) -> None:
        if self._running:
            try:
                with self._lock:
                    self._cam.set_controls(ctrl)
            except Exception as exc:
                log.warning("apply_controls error: %s", exc)

    def stop(self) -> None:
        if self._running:
            try:
                self._cam.stop()
                self._running = False
                log.info("Camera: stopped")
            except Exception as exc:
                log.warning("Camera stop error: %s", exc)

    def close(self) -> None:
        self.stop()
        if self._cam:
            try:
                self._cam.close()
                log.info("Camera: closed")
            except Exception as exc:
                log.warning("Camera close error: %s", exc)


# ══════════════════════════════════════════════════════════════════════════════
# TimelapseController
# ══════════════════════════════════════════════════════════════════════════════
class TimelapseController:
    """
    Manages timelapse capture in a daemon background thread.
    Callbacks (on_frame, on_finish, on_error) are called from that thread;
    the owner must schedule any tkinter updates via root.after(0, fn).
    """

    def __init__(self, camera: CameraController):
        self._cam         = camera
        self._stop_event  = threading.Event()
        self._thread      = None
        self._count       = 0

        self.on_frame     = None   # callable(frame_idx: int, filepath: str)
        self.on_finish    = None   # callable(total: int)
        self.on_error     = None   # callable(msg: str)
        self.on_light_on  = None   # callable() – called on the worker thread after light turns on
        self.on_light_off = None   # callable() – called on the worker thread after light turns off
        self.light        = None   # LightController – set by owner; if set, light flashes per frame
        self.warmup_secs  = 1.0    # seconds to wait after turning light on before capturing

    def start(self, interval_secs: float, total_frames: int,
              prefix: str, save_dir: str) -> None:
        if self.is_running():
            return
        os.makedirs(save_dir, exist_ok=True)
        self._stop_event.clear()
        self._count = 0
        self._thread = threading.Thread(
            target=self._run,
            args=(interval_secs, total_frames, prefix, save_dir),
            daemon=True,
        )
        self._thread.start()
        log.info("Timelapse started: interval=%.1fs frames=%s",
                 interval_secs, total_frames or "∞")

    def stop(self) -> None:
        self._stop_event.set()
        log.info("Timelapse stop requested")

    def is_running(self) -> bool:
        return self._thread is not None and self._thread.is_alive()

    def frame_count(self) -> int:
        return self._count

    def _run(self, interval_secs, total_frames, prefix, save_dir):
        while not self._stop_event.is_set():
            # Turn on light and wait for warmup before capturing
            if self.light is not None:
                self.light.turn_on()
                if self.on_light_on:
                    self.on_light_on()
                if self._stop_event.wait(timeout=self.warmup_secs):
                    self.light.turn_off()
                    if self.on_light_off:
                        self.on_light_off()
                    break

            ts = datetime.now().strftime("%Y%m%d_%H%M%S")
            self._count += 1
            filename = f"{prefix}_{ts}_{self._count:05d}.jpg"
            filepath = os.path.join(save_dir, filename)

            ok = self._cam.capture_timelapse_frame(filepath)

            # Turn off light immediately after capture
            if self.light is not None:
                self.light.turn_off()
                if self.on_light_off:
                    self.on_light_off()

            if ok:
                if self.on_frame:
                    self.on_frame(self._count, filepath)
            else:
                if self.on_error:
                    self.on_error(f"Frame {self._count} failed to capture")

            if total_frames > 0 and self._count >= total_frames:
                break

            # Interruptible sleep – wakes immediately when stop() is called
            self._stop_event.wait(timeout=interval_secs)

        total = self._count
        if self.on_finish:
            self.on_finish(total)
        log.info("Timelapse finished: %d frames", total)


# ══════════════════════════════════════════════════════════════════════════════
# ScrollableFrame – reusable scrollable container
# ══════════════════════════════════════════════════════════════════════════════
class ScrollableFrame(tk.Frame):
    """A tk.Frame with a vertically scrollable inner frame (self.inner)."""

    def __init__(self, parent, bg=C["bg"], **kw):
        super().__init__(parent, bg=bg, **kw)

        self._canvas = tk.Canvas(self, bg=bg, highlightthickness=0)
        scrollbar = tk.Scrollbar(self, orient="vertical",
                                  command=self._canvas.yview)
        self.inner = tk.Frame(self._canvas, bg=bg)

        self._win_id = self._canvas.create_window(
            (0, 0), window=self.inner, anchor="nw"
        )

        self._canvas.configure(yscrollcommand=scrollbar.set)
        self._canvas.pack(side="left", fill="both", expand=True)
        scrollbar.pack(side="right", fill="y")

        self.inner.bind("<Configure>", self._on_inner_configure)
        self._canvas.bind("<Configure>", self._on_canvas_configure)

    def activate_scroll(self):
        """Claim the global mousewheel binding for this canvas.
        Call whenever this frame's tab becomes visible."""
        self._canvas.bind_all("<MouseWheel>",
                               lambda e: self._canvas.yview_scroll(
                                   -1 * (e.delta // 120), "units"))
        self._canvas.bind_all("<Button-4>",
                               lambda e: self._canvas.yview_scroll(-1, "units"))
        self._canvas.bind_all("<Button-5>",
                               lambda e: self._canvas.yview_scroll(1, "units"))

    def _on_inner_configure(self, _event):
        self._canvas.configure(scrollregion=self._canvas.bbox("all"))

    def _on_canvas_configure(self, event):
        self._canvas.itemconfig(self._win_id, width=event.width)


# ══════════════════════════════════════════════════════════════════════════════
# dishcamApp – main application window
# ══════════════════════════════════════════════════════════════════════════════
class dishcamApp(tk.Tk):
    """
    Root tkinter window.  Layout (top → bottom):
        Header    90 px   – title, status, close button
        LightBar  75 px   – always-visible brightness slider
        Preview  480 px   – live camera feed
        TabBar    70 px   – Capture | Timelapse | Settings | About
        Content  565 px   – active tab
    Total: 1280 px
    """

    # Fixed-height chrome: header(90) + lightbar(75) + prev-brightness(58) + tabbar(70)
    _CHROME_H    = 293
    _MIN_CONTENT = 200   # minimum pixels for the scrollable tab area
    _MIN_PREVIEW = 40    # preview can get very short; scrollable content is what matters

    def __init__(self):
        super().__init__()
        self.title("Dishcam")
        self.configure(bg=C["bg"])

        # Adapt to actual screen height so nothing falls off the bottom.
        screen_h        = self.winfo_screenheight()
        win_h           = min(SCREEN_H, screen_h - 50)   # leave ~50 px for taskbar
        self._preview_h = max(
            self._MIN_PREVIEW,
            win_h - self._CHROME_H - self._MIN_CONTENT,
        )

        self.geometry(f"{SCREEN_W}x{win_h}+0+0")
        self.resizable(False, True)   # fixed width, variable height
        # Uncomment for true kiosk mode:
        # self.attributes("-fullscreen", True)

        # ── Controllers ───────────────────────────────────────────────────────
        self._light     = LightController(prefer_sysfs=True)
        self._camera    = CameraController()
        self._timelapse = TimelapseController(self._camera)

        self._timelapse.on_frame     = self._tl_on_frame
        self._timelapse.on_finish    = self._tl_on_finish
        self._timelapse.on_error     = self._tl_on_error
        self._timelapse.on_light_on  = self._tl_on_light_on
        self._timelapse.on_light_off = self._tl_on_light_off
        self._timelapse.light        = self._light

        # ── Preview state ─────────────────────────────────────────────────────
        self._preview_queue = queue.Queue(maxsize=1)
        self._preview_stop  = threading.Event()
        self._imgtk_ref     = None   # keep PhotoImage alive

        # ── tk variables ──────────────────────────────────────────────────────
        self._light_pct    = tk.DoubleVar(value=0.0)
        self._save_dir     = tk.StringVar(value=DEFAULT_SAVE_DIR)
        self._cap_prefix   = tk.StringVar(value="capture")
        self._tl_prefix    = tk.StringVar(value="timelapse")
        self._tl_save_dir        = tk.StringVar(value=DEFAULT_SAVE_DIR)
        self._preview_brightness = tk.DoubleVar(value=1.0)
        self._pre_tl_brightness  = 0.0
        self._tl_hours     = tk.IntVar(value=0)
        self._tl_minutes   = tk.IntVar(value=5)
        self._tl_seconds   = tk.IntVar(value=0)
        self._tl_total     = tk.IntVar(value=0)
        self._ae_var       = tk.BooleanVar(value=True)
        self._awb_var      = tk.BooleanVar(value=True)
        self._exp_var      = tk.StringVar(value=str(DEFAULT_EXPOSURE_US))
        self._gain_var     = tk.DoubleVar(value=DEFAULT_GAIN)
        self._pwm_freq_var = tk.StringVar(value=str(PWM_FREQ_HZ))
        self._pwm_hw_var   = tk.BooleanVar(value=True)
        self._status_text  = tk.StringVar(value="Ready")

        # ── Build UI ──────────────────────────────────────────────────────────
        self._tab_frames   = {}
        self._tab_buttons  = {}

        self._build_header()
        self._build_lightbar()
        self._build_preview_brightness_bar()
        self._build_preview()
        self._build_tabbar()
        self._build_content()

        self.protocol("WM_DELETE_WINDOW", self._on_close)

        # ── Start preview loop (shows "Initialising..." until camera is ready) ─
        self._start_preview_thread()
        self._schedule_preview()

        # ── Camera starts in background – GUI is never blocked ────────────────
        self._status_text.set("Initialising camera...")
        threading.Thread(target=self._init_camera_bg, daemon=True).start()

    # ══════════════════════════════════════════════════════════════════════════
    # UI construction
    # ══════════════════════════════════════════════════════════════════════════

    def _build_header(self):
        hdr = tk.Frame(self, bg=C["panel"], height=90)
        hdr.pack(fill="x", side="top")
        hdr.pack_propagate(False)

        tk.Label(hdr, text="Dishcam", font=(FF, 30, "bold"),
                 bg=C["panel"], fg=C["orange"]).pack(side="left", padx=20)

        # Close / quit button
        tk.Button(hdr, text="✕", font=(FF, 22, "bold"),
                  bg=C["panel"], fg=C["mgray"],
                  activebackground=C["red"], activeforeground=C["white"],
                  relief="flat", padx=16, pady=0,
                  command=self._on_close).pack(side="right", padx=10)

        # Status dots – camera starts amber (initialising), turns green/red later
        sf = tk.Frame(hdr, bg=C["panel"])
        sf.pack(side="right", padx=6)
        self._cam_dot = tk.Label(sf, text="●", font=(FF, 20),
                                  bg=C["panel"], fg="#FFAA00")   # amber = initialising
        self._cam_dot.grid(row=0, column=0)
        tk.Label(sf, text="CAM", font=(FF, 14),
                 bg=C["panel"], fg=C["mgray"]).grid(row=0, column=1, padx=(3, 14))
        light_ok = self._light.backend != "none"
        self._light_dot = tk.Label(sf, text="●", font=(FF, 20),
                 bg=C["panel"],
                 fg=C["mgray"] if light_ok else C["red"])
        self._light_dot.grid(row=0, column=2)
        tk.Label(sf, text="LIGHT", font=(FF, 14),
                 bg=C["panel"], fg=C["mgray"]).grid(row=0, column=3, padx=(3, 0))

        # Status message
        tk.Label(hdr, textvariable=self._status_text, font=(FF, 15),
                 bg=C["panel"], fg=C["lgray"]).pack(side="left", padx=16)

    def _build_lightbar(self):
        bar = tk.Frame(self, bg=C["card"], height=75)
        bar.pack(fill="x")
        bar.pack_propagate(False)

        tk.Label(bar, text="Light", font=(FF, 17, "bold"),
                 bg=C["card"], fg=C["white"]).pack(side="left", padx=(18, 6))

        self._light_pct_lbl = tk.Label(bar, text="0%", width=5,
                                        font=(FF, 17, "bold"),
                                        bg=C["card"], fg=C["orange"])
        self._light_pct_lbl.pack(side="left", padx=4)

        slider = tk.Scale(bar, from_=0, to=100,
                          orient="horizontal", variable=self._light_pct,
                          command=self._on_light_slider,
                          resolution=5,          # snap to 5 % increments
                          bg=C["card"], fg=C["orange"],
                          troughcolor=C["dgray"],
                          highlightthickness=0,
                          activebackground=C["orhi"],
                          sliderlength=52, width=26,
                          showvalue=False)
        slider.pack(side="left", fill="x", expand=True, padx=6)

        tk.Button(bar, text="ON", font=(FF, 16, "bold"),
                  bg=C["orange"], fg=C["white"],
                  activebackground=C["orhi"], relief="flat",
                  padx=14, pady=10,
                  command=lambda: self._set_light(100)).pack(side="right", padx=(6, 18))
        tk.Button(bar, text="OFF", font=(FF, 16, "bold"),
                  bg=C["dgray"], fg=C["white"],
                  activebackground=C["mgray"], relief="flat",
                  padx=14, pady=10,
                  command=lambda: self._set_light(0)).pack(side="right", padx=6)

    def _build_preview_brightness_bar(self):
        bar = tk.Frame(self, bg=C["bg"], height=58)
        bar.pack(fill="x")
        bar.pack_propagate(False)

        tk.Label(bar, text="Preview", font=(FF, 14, "bold"),
                 bg=C["bg"], fg=C["mgray"]).pack(side="left", padx=(18, 6))

        self._prev_bright_lbl = tk.Label(bar, text="1.0×", width=5,
                                          font=(FF, 14, "bold"),
                                          bg=C["bg"], fg=C["lgray"])
        self._prev_bright_lbl.pack(side="left", padx=4)

        def _on_change(v):
            self._prev_bright_lbl.configure(text=f"{float(v):.1f}×")

        tk.Scale(bar, from_=0.1, to=3.0,
                 orient="horizontal", variable=self._preview_brightness,
                 command=_on_change,
                 resolution=0.1,
                 bg=C["bg"], fg=C["lgray"],
                 troughcolor=C["dgray"],
                 highlightthickness=0,
                 activebackground=C["orhi"],
                 sliderlength=42, width=20,
                 showvalue=False).pack(side="left", fill="x", expand=True, padx=6)

        def _reset():
            self._preview_brightness.set(1.0)
            self._prev_bright_lbl.configure(text="1.0×")

        tk.Button(bar, text="1×", font=(FF, 13),
                  bg=C["dgray"], fg=C["white"],
                  activebackground=C["mgray"], relief="flat",
                  padx=10, pady=5,
                  command=_reset).pack(side="right", padx=(6, 18))

    def _build_preview(self):
        pf = tk.Frame(self, bg="black", height=self._preview_h)
        pf.pack(fill="x")
        pf.pack_propagate(False)

        self._preview_lbl = tk.Label(pf, bg="black",
                                      text="Initialising camera...",
                                      fg=C["mgray"], font=(FF, 18))
        self._preview_lbl.pack(fill="both", expand=True)

    def _build_tabbar(self):
        bar = tk.Frame(self, bg=C["panel"], height=70)
        bar.pack(fill="x")
        bar.pack_propagate(False)

        tabs = [
            ("capture",   "Capture"),
            ("timelapse", "Timelapse"),
            ("settings",  "Settings"),
            ("about",     "About"),
        ]
        for name, label in tabs:
            btn = tk.Button(bar, text=label,
                            font=(FF, 18, "bold"),
                            bg=C["orange"] if name == "capture" else C["panel"],
                            fg=C["white"] if name == "capture" else C["mgray"],
                            activebackground=C["orhi"],
                            activeforeground=C["white"],
                            relief="flat",
                            command=lambda n=name: self._switch_tab(n))
            btn.pack(side="left", fill="both", expand=True)
            self._tab_buttons[name] = btn

    def _build_content(self):
        container = tk.Frame(self, bg=C["bg"])
        container.pack(fill="both", expand=True)

        for name, builder in [
            ("capture",   self._build_capture_tab),
            ("timelapse", self._build_timelapse_tab),
            ("settings",  self._build_settings_tab),
            ("about",     self._build_about_tab),
        ]:
            f = builder(container)
            self._tab_frames[name] = f

        self._switch_tab("capture")

    def _switch_tab(self, name: str):
        for n, f in self._tab_frames.items():
            if n == name:
                f.pack(fill="both", expand=True)
                if isinstance(f, ScrollableFrame):
                    f.activate_scroll()
            else:
                f.pack_forget()
        for n, btn in self._tab_buttons.items():
            if n == name:
                btn.configure(bg=C["orange"], fg=C["white"])
            else:
                btn.configure(bg=C["panel"], fg=C["mgray"])

    # ══════════════════════════════════════════════════════════════════════════
    # Capture tab
    # ══════════════════════════════════════════════════════════════════════════

    def _build_capture_tab(self, parent) -> tk.Frame:
        sf = ScrollableFrame(parent, bg=C["bg"])
        frame = sf.inner

        # Save folder
        fc = tk.Frame(frame, bg=C["card"])
        fc.pack(fill="x", padx=16, pady=(16, 0))
        tk.Label(fc, text="Save folder", font=(FF, 16),
                 bg=C["card"], fg=C["mgray"]).pack(anchor="w", padx=16, pady=(10, 0))
        dr = tk.Frame(fc, bg=C["card"])
        dr.pack(fill="x", padx=16, pady=(6, 12))
        tk.Entry(dr, textvariable=self._save_dir, font=(FF, 17),
                 bg=C["bg"], fg=C["white"], insertbackground=C["orange"],
                 relief="flat").pack(side="left", fill="x", expand=True, ipady=11)
        tk.Button(dr, text="Browse", font=(FF, 16),
                  bg=C["orlo"], fg=C["white"],
                  activebackground=C["orange"], relief="flat",
                  padx=14, pady=6,
                  command=self._pick_folder).pack(side="right", padx=(10, 0))

        # Prefix
        pc = tk.Frame(frame, bg=C["card"])
        pc.pack(fill="x", padx=16, pady=10)
        tk.Label(pc, text="Filename prefix", font=(FF, 16),
                 bg=C["card"], fg=C["mgray"]).pack(anchor="w", padx=16, pady=(10, 0))
        tk.Entry(pc, textvariable=self._cap_prefix, font=(FF, 20),
                 bg=C["bg"], fg=C["white"], insertbackground=C["orange"],
                 relief="flat").pack(fill="x", padx=16, pady=(6, 12), ipady=13)

        # Capture button
        self._cap_btn = tk.Button(
            frame, text="CAPTURE PHOTO",
            font=(FF, 30, "bold"),
            bg=C["orange"], fg=C["white"],
            activebackground=C["orhi"], activeforeground=C["white"],
            relief="flat", pady=30,
            command=self._do_capture,
        )
        self._cap_btn.pack(fill="x", padx=16, pady=14)

        # Status
        self._cap_status = tk.Label(frame, text="Ready",
                                     font=(FF, 17), wraplength=680,
                                     bg=C["bg"], fg=C["lgray"])
        self._cap_status.pack(padx=16, pady=4)

        return sf

    # ══════════════════════════════════════════════════════════════════════════
    # Timelapse tab
    # ══════════════════════════════════════════════════════════════════════════

    def _build_timelapse_tab(self, parent) -> tk.Frame:
        sf = ScrollableFrame(parent, bg=C["bg"])
        frame = sf.inner

        # Interval H:M:S
        ic = tk.Frame(frame, bg=C["card"])
        ic.pack(fill="x", padx=16, pady=(16, 0))
        tk.Label(ic, text="Interval between captures", font=(FF, 16),
                 bg=C["card"], fg=C["mgray"]).pack(anchor="w", padx=16, pady=(10, 6))
        ir = tk.Frame(ic, bg=C["card"])
        ir.pack(fill="x", padx=16, pady=(0, 12))
        for var, label, maxv in [
            (self._tl_hours,   "h", 23),
            (self._tl_minutes, "m", 59),
            (self._tl_seconds, "s", 59),
        ]:
            sb_f = tk.Frame(ir, bg=C["card"])
            sb_f.pack(side="left", padx=8)
            sb = tk.Spinbox(sb_f, from_=0, to=maxv, textvariable=var,
                            width=4, font=(FF, 24),
                            bg=C["bg"], fg=C["white"],
                            buttonbackground=C["dgray"],
                            insertbackground=C["orange"],
                            relief="flat", wrap=True)
            sb.pack(side="left", ipady=10)
            tk.Label(sb_f, text=label, font=(FF, 20),
                     bg=C["card"], fg=C["lgray"]).pack(side="left", padx=6)

        # Total frames
        nc = tk.Frame(frame, bg=C["card"])
        nc.pack(fill="x", padx=16, pady=10)
        tk.Label(nc, text="Total frames  (0 = unlimited)", font=(FF, 16),
                 bg=C["card"], fg=C["mgray"]).pack(anchor="w", padx=16, pady=(10, 6))
        tk.Spinbox(nc, from_=0, to=99999, textvariable=self._tl_total,
                   width=7, font=(FF, 24),
                   bg=C["bg"], fg=C["white"],
                   buttonbackground=C["dgray"],
                   insertbackground=C["orange"],
                   relief="flat").pack(anchor="w", padx=16, pady=(0, 12), ipady=10)

        # Prefix
        prc = tk.Frame(frame, bg=C["card"])
        prc.pack(fill="x", padx=16, pady=0)
        tk.Label(prc, text="Filename prefix", font=(FF, 16),
                 bg=C["card"], fg=C["mgray"]).pack(anchor="w", padx=16, pady=(10, 0))
        tk.Entry(prc, textvariable=self._tl_prefix, font=(FF, 20),
                 bg=C["bg"], fg=C["white"], insertbackground=C["orange"],
                 relief="flat").pack(fill="x", padx=16, pady=(6, 12), ipady=13)

        # Save folder
        fc = tk.Frame(frame, bg=C["card"])
        fc.pack(fill="x", padx=16, pady=10)
        tk.Label(fc, text="Save folder", font=(FF, 16),
                 bg=C["card"], fg=C["mgray"]).pack(anchor="w", padx=16, pady=(10, 0))
        dr = tk.Frame(fc, bg=C["card"])
        dr.pack(fill="x", padx=16, pady=(6, 12))
        tk.Entry(dr, textvariable=self._tl_save_dir, font=(FF, 17),
                 bg=C["bg"], fg=C["white"], insertbackground=C["orange"],
                 relief="flat").pack(side="left", fill="x", expand=True, ipady=11)
        tk.Button(dr, text="Browse", font=(FF, 16),
                  bg=C["orlo"], fg=C["white"],
                  activebackground=C["orange"], relief="flat",
                  padx=14, pady=6,
                  command=self._pick_tl_folder).pack(side="right", padx=(10, 0))

        # Start / Stop
        btn_row = tk.Frame(frame, bg=C["bg"])
        btn_row.pack(fill="x", padx=16, pady=14)
        self._tl_start_btn = tk.Button(
            btn_row, text="▶  START",
            font=(FF, 24, "bold"), bg=C["orange"], fg=C["white"],
            activebackground=C["orhi"], relief="flat",
            pady=20, command=self._tl_start,
        )
        self._tl_start_btn.pack(side="left", fill="x", expand=True, padx=(0, 8))

        self._tl_stop_btn = tk.Button(
            btn_row, text="■  STOP",
            font=(FF, 24, "bold"), bg=C["dgray"], fg=C["white"],
            activebackground=C["red"], relief="flat",
            pady=20, state="disabled",
            command=self._tl_stop,
        )
        self._tl_stop_btn.pack(side="right", fill="x", expand=True, padx=(8, 0))

        # Progress
        self._tl_progress = tk.Label(frame, text="Not running",
                                      font=(FF, 17), bg=C["bg"], fg=C["lgray"])
        self._tl_progress.pack(padx=16, pady=(6, 0))
        self._tl_last_file = tk.Label(frame, text="",
                                       font=(FF, 15), bg=C["bg"], fg=C["mgray"],
                                       wraplength=680)
        self._tl_last_file.pack(padx=16, pady=2)

        return sf

    # ══════════════════════════════════════════════════════════════════════════
    # Settings tab
    # ══════════════════════════════════════════════════════════════════════════

    def _build_settings_tab(self, parent) -> tk.Frame:
        sf = ScrollableFrame(parent, bg=C["bg"])
        inner = sf.inner

        # ── Section: Presets ──────────────────────────────────────────────────
        self._section_label(inner, "Presets")

        preset_card = tk.Frame(inner, bg=C["card"])
        preset_card.pack(fill="x", padx=16, pady=4)
        tk.Label(preset_card,
                 text="Fixed macro / timelapse preset\n"
                      "(AE off, AWB off, 1 s exposure, gain 1.0)",
                 font=(FF, 15), bg=C["card"], fg=C["lgray"],
                 justify="left").pack(side="left", padx=16, pady=14)
        tk.Button(preset_card, text="Load Preset",
                  font=(FF, 16, "bold"),
                  bg=C["orlo"], fg=C["white"],
                  activebackground=C["orange"], relief="flat",
                  padx=16, pady=10,
                  command=self._load_fixedlight_preset,
                  ).pack(side="right", padx=16, pady=10)

        # ── Section: Exposure ─────────────────────────────────────────────────
        self._section_label(inner, "Exposure")

        self._ae_btn = self._toggle_row(
            inner, "Auto Exposure (AE)", self._ae_var, self._on_ae_toggle)

        # Exposure time
        self._exp_card = tk.Frame(inner, bg=C["card"])
        self._exp_card.pack(fill="x", padx=16, pady=4)
        tk.Label(self._exp_card, text="Exposure time (µs)", font=(FF, 17),
                 bg=C["card"], fg=C["white"]).pack(anchor="w", padx=16, pady=(12, 0))
        exp_r = tk.Frame(self._exp_card, bg=C["card"])
        exp_r.pack(fill="x", padx=16, pady=(6, 12))
        self._exp_entry = tk.Entry(
            exp_r, textvariable=self._exp_var, font=(FF, 20),
            bg=C["bg"], fg=C["white"], insertbackground=C["orange"],
            relief="flat", state="disabled")
        self._exp_entry.pack(side="left", fill="x", expand=True, ipady=13)
        tk.Label(exp_r, text="µs", font=(FF, 16), bg=C["card"],
                 fg=C["mgray"]).pack(side="right", padx=10)

        # Analogue gain slider
        self._slider_row(inner, "Analogue Gain", self._gain_var,
                         1.0, 16.0, 0.1, fmt="{:.1f}×")

        # ── Section: White Balance ────────────────────────────────────────────
        self._section_label(inner, "White Balance")
        self._toggle_row(inner, "Auto White Balance (AWB)",
                         self._awb_var, lambda: None)

        # ── Apply / Reset camera buttons ──────────────────────────────────────
        tk.Button(inner, text="Apply Camera Settings",
                  font=(FF, 20, "bold"),
                  bg=C["orange"], fg=C["white"],
                  activebackground=C["orhi"], relief="flat",
                  pady=20, command=self._apply_cam_settings,
                  ).pack(fill="x", padx=16, pady=(20, 4))
        tk.Button(inner, text="Reset to Default (Auto)",
                  font=(FF, 16, "bold"),
                  bg=C["dgray"], fg=C["white"],
                  activebackground=C["mgray"], relief="flat",
                  pady=12, command=self._reset_cam_settings,
                  ).pack(fill="x", padx=16, pady=(0, 4))

        self._settings_status = tk.Label(inner, text="",
                                          font=(FF, 16), bg=C["bg"], fg=C["green"])
        self._settings_status.pack(padx=16)

        # ── Section: Light / PWM ──────────────────────────────────────────────
        self._section_label(inner, "Light / PWM")

        self._toggle_row(inner, "Hardware PWM (sysfs)",
                         self._pwm_hw_var, self._reinit_light)
        tk.Label(inner,
                 text="⚠  Switching to software PWM disables hardware PWM\n"
                      "    until the next reboot.",
                 font=(FF, 13), bg=C["bg"], fg=C["amber"],
                 justify="left").pack(anchor="w", padx=32, pady=(0, 6))

        freq_card = tk.Frame(inner, bg=C["card"])
        freq_card.pack(fill="x", padx=16, pady=4)
        tk.Label(freq_card, text="PWM frequency (Hz)", font=(FF, 17),
                 bg=C["card"], fg=C["white"]).pack(anchor="w", padx=16, pady=(12, 0))
        tk.Label(freq_card,
                 text="Lower = visible flicker but more MOSFET-friendly.\n"
                      "Higher = smoother but may cause heating on some MOSFETs.\n"
                      "1000 Hz is a safe default.",
                 font=(FF, 12), bg=C["card"], fg=C["mgray"],
                 justify="left").pack(anchor="w", padx=16, pady=(2, 0))
        freq_row = tk.Frame(freq_card, bg=C["card"])
        freq_row.pack(fill="x", padx=16, pady=(6, 12))
        freq_spinbox = tk.Spinbox(
            freq_row,
            values=(100, 200, 500, 1000, 2000, 5000, 10000, 20000),
            textvariable=self._pwm_freq_var,
            font=(FF, 20), width=8,
            bg=C["bg"], fg=C["white"],
            buttonbackground=C["dgray"],
            insertbackground=C["orange"],
            relief="flat")
        freq_spinbox.pack(side="left", ipady=10)
        tk.Label(freq_row, text="Hz", font=(FF, 16),
                 bg=C["card"], fg=C["mgray"]).pack(side="left", padx=10)
        tk.Button(freq_row, text="Apply Freq",
                  font=(FF, 15, "bold"),
                  bg=C["orlo"], fg=C["white"],
                  activebackground=C["orange"], relief="flat",
                  padx=14, pady=8,
                  command=self._apply_pwm_freq).pack(side="right")

        self._pwm_freq_status = tk.Label(inner, text="",
                                          font=(FF, 14), bg=C["bg"], fg=C["green"])
        self._pwm_freq_status.pack(padx=16, pady=(0, 16))

        return sf

    # ── settings helpers ──────────────────────────────────────────────────────

    def _section_label(self, parent, text: str):
        tk.Label(parent, text=text,
                 font=(FF, 18, "bold"),
                 bg=C["bg"], fg=C["orange"]).pack(
                     anchor="w", padx=16, pady=(18, 2))

    def _toggle_row(self, parent, label: str, var: tk.BooleanVar,
                    on_change) -> tk.Button:
        row = tk.Frame(parent, bg=C["card"])
        row.pack(fill="x", padx=16, pady=4)
        tk.Label(row, text=label, font=(FF, 17),
                 bg=C["card"], fg=C["white"]).pack(side="left", padx=16, pady=16)
        btn = tk.Button(row, font=(FF, 16, "bold"),
                        fg=C["white"], relief="flat", padx=24, pady=10)
        btn.pack(side="right", padx=16, pady=10)

        def _refresh(*_):
            btn.configure(text="ON" if var.get() else "OFF",
                          bg=C["green"] if var.get() else C["dgray"])

        def _toggle():
            var.set(not var.get())
            _refresh()
            on_change()

        btn.configure(command=_toggle)
        _refresh()
        return btn

    def _slider_row(self, parent, label: str, var: tk.DoubleVar,
                    from_: float, to: float, resolution: float,
                    fmt: str = "{:.1f}"):
        card = tk.Frame(parent, bg=C["card"])
        card.pack(fill="x", padx=16, pady=4)
        hdr = tk.Frame(card, bg=C["card"])
        hdr.pack(fill="x", padx=16, pady=(12, 0))
        tk.Label(hdr, text=label, font=(FF, 17),
                 bg=C["card"], fg=C["white"]).pack(side="left")
        val_lbl = tk.Label(hdr, text=fmt.format(var.get()),
                            font=(FF, 17), bg=C["card"], fg=C["orange"])
        val_lbl.pack(side="right")

        def _cb(v):
            val_lbl.configure(text=fmt.format(float(v)))

        tk.Scale(card, from_=from_, to=to, resolution=resolution,
                 orient="horizontal", variable=var, command=_cb,
                 bg=C["card"], fg=C["orange"],
                 troughcolor=C["dgray"], highlightthickness=0,
                 activebackground=C["orhi"],
                 sliderlength=52, width=28, showvalue=False,
                 ).pack(fill="x", padx=16, pady=(6, 12))

    # ══════════════════════════════════════════════════════════════════════════
    # About tab
    # ══════════════════════════════════════════════════════════════════════════

    def _build_about_tab(self, parent) -> tk.Frame:
        sf = ScrollableFrame(parent, bg=C["bg"])
        frame = sf.inner

        def lbl(text, **kw):
            tk.Label(frame, text=text, bg=C["bg"],
                     wraplength=680, justify="left", **kw).pack(
                         anchor="w", padx=28, pady=3)

        lbl(f"Dishcam  v{APP_VERSION}",
            font=(FF, 26, "bold"), fg=C["orange"])
        lbl("Petri dish imaging for Raspberry Pi",
            font=(FF, 17), fg=C["lgray"])

        tk.Frame(frame, bg=C["border"], height=1).pack(fill="x", padx=16, pady=14)

        lbl("GPIO / Light", font=(FF, 17, "bold"), fg=C["white"])
        lbl(f"  Light pin : GPIO {GPIO_LIGHT} (BCM) – physical pin 12",
            font=(FF, 16), fg=C["lgray"])
        lbl(f"  PWM backend: {self._light.backend}",
            font=(FF, 16), fg=C["lgray"])
        lbl(f"  PWM freq   : {PWM_FREQ_HZ} Hz",
            font=(FF, 16), fg=C["lgray"])
        lbl("  Connect LED/MOSFET gate to GPIO 18, GND to GND.",
            font=(FF, 15), fg=C["mgray"])

        tk.Frame(frame, bg=C["border"], height=1).pack(fill="x", padx=16, pady=14)

        lbl("Camera", font=(FF, 17, "bold"), fg=C["white"])
        self._about_cam_var = tk.StringVar(value="  Initialising...")
        tk.Label(frame, textvariable=self._about_cam_var,
                 bg=C["bg"], fg=C["lgray"],
                 font=(FF, 16), justify="left").pack(anchor="w", padx=28, pady=3)

        tk.Frame(frame, bg=C["border"], height=1).pack(fill="x", padx=16, pady=14)

        lbl("Libraries: picamera2 · RPi.GPIO · Pillow · tkinter",
            font=(FF, 15), fg=C["mgray"])

        return sf

    # ══════════════════════════════════════════════════════════════════════════
    # Preview thread & update loop
    # ══════════════════════════════════════════════════════════════════════════

    def _init_camera_bg(self):
        """Run camera start() in a background thread; update UI when done."""
        self._camera.start()
        def _on_done():
            if self._camera.available:
                self._cam_dot.configure(fg=C["green"])
                self._status_text.set("Camera ready")
                cam = self._camera._cam
                try:
                    model  = cam.camera_properties.get("Model", "unknown")
                    sensor = cam.sensor_resolution
                    self._about_cam_var.set(
                        f"  Model  : {model}\n  Sensor : {sensor[0]}×{sensor[1]}")
                except Exception:
                    self._about_cam_var.set("  Camera ready")
            else:
                self._cam_dot.configure(fg=C["red"])
                self._status_text.set("Camera failed – check connection")
                self._about_cam_var.set("  Camera failed to start")
        self.after(0, _on_done)

    def _start_preview_thread(self):
        self._preview_stop.clear()
        t = threading.Thread(target=self._preview_worker, daemon=True)
        t.start()

    def _preview_worker(self):
        """Background thread: capture frames and push to queue."""
        while not self._preview_stop.is_set():
            if not PIL_AVAILABLE:
                time.sleep(0.5)
                continue
            arr = self._camera.capture_frame()
            if arr is not None:
                try:
                    img = Image.fromarray(arr)
                    self._preview_queue.put_nowait(img)
                except queue.Full:
                    pass   # drop stale frame
                except Exception as exc:
                    log.debug("Preview worker error: %s", exc)
            if arr is None:
                time.sleep(0.1)

    def _schedule_preview(self):
        self._update_preview()

    def _update_preview(self):
        try:
            img = self._preview_queue.get_nowait()
            brightness = self._preview_brightness.get()
            if abs(brightness - 1.0) > 0.05:
                img = ImageEnhance.Brightness(img).enhance(brightness)
            imgtk = ImageTk.PhotoImage(image=img)
            self._preview_lbl.imgtk = imgtk      # prevent GC
            self._preview_lbl.configure(image=imgtk, text="")
        except queue.Empty:
            pass
        except Exception as exc:
            log.debug("Preview update error: %s", exc)
        finally:
            self.after(1000 // PREVIEW_FPS, self._update_preview)

    # ══════════════════════════════════════════════════════════════════════════
    # Light actions
    # ══════════════════════════════════════════════════════════════════════════

    def _on_light_slider(self, value: str):
        pct = float(value)
        self._light_pct_lbl.configure(text=f"{pct:.0f}%")
        if pct > 0:
            self._light.turn_on()
            self._light.set_brightness(pct)
        else:
            self._light.turn_off()
        self._update_light_dot()

    def _update_light_dot(self):
        if self._light.backend == "none":
            return
        self._light_dot.configure(fg=C["orange"] if self._light.is_on() else C["mgray"])

    def _set_light(self, pct: float):
        self._light_pct.set(pct)
        self._on_light_slider(str(pct))

    # ══════════════════════════════════════════════════════════════════════════
    # Capture actions
    # ══════════════════════════════════════════════════════════════════════════

    def _pick_folder(self):
        d = filedialog.askdirectory(initialdir=self._save_dir.get())
        if d:
            self._save_dir.set(d)

    def _pick_tl_folder(self):
        d = filedialog.askdirectory(initialdir=self._tl_save_dir.get())
        if d:
            self._tl_save_dir.set(d)

    def _do_capture(self):
        out_dir  = self._save_dir.get()
        prefix   = self._cap_prefix.get().strip() or "capture"
        ts       = datetime.now().strftime("%Y%m%d_%H%M%S")
        filepath = os.path.join(out_dir, f"{prefix}_{ts}.jpg")

        os.makedirs(out_dir, exist_ok=True)

        self._cap_btn.configure(state="disabled", bg=C["orlo"])
        self._cap_status.configure(text="Capturing...", fg=C["lgray"])
        self._status_text.set("Capturing...")

        def on_done(ok: bool, fp: str):
            def _update():
                self._cap_btn.configure(state="normal", bg=C["orange"])
                if ok:
                    self._cap_status.configure(
                        text=f"Saved: {os.path.basename(fp)}", fg=C["green"])
                    self._status_text.set("Captured OK")
                else:
                    self._cap_status.configure(text="Capture failed", fg=C["red"])
                    self._status_text.set("Capture failed")
            self.after(0, _update)

        if self._camera.available:
            self._camera.capture_still(filepath, on_done)
        else:
            def _demo():
                time.sleep(1)
                on_done(True, filepath)
            threading.Thread(target=_demo, daemon=True).start()

    # ══════════════════════════════════════════════════════════════════════════
    # Timelapse actions
    # ══════════════════════════════════════════════════════════════════════════

    def _tl_start(self):
        h = self._tl_hours.get()
        m = self._tl_minutes.get()
        s = self._tl_seconds.get()
        interval = h * 3600 + m * 60 + s
        if interval < 1:
            messagebox.showerror("Invalid interval",
                                  "Please set an interval of at least 1 second.")
            return

        prefix   = self._tl_prefix.get().strip() or "timelapse"
        total    = self._tl_total.get()
        save_dir = self._tl_save_dir.get()

        self._pre_tl_brightness = self._light_pct.get()
        self._timelapse.start(float(interval), total, prefix, save_dir)

        self._tl_start_btn.configure(state="disabled", bg=C["orlo"])
        self._tl_stop_btn.configure(state="normal", bg=C["red"])
        self._tl_progress.configure(
            text=f"Running...  0 / {'∞' if total == 0 else total} frames",
            fg=C["green"])
        self._tl_last_file.configure(text="")
        self._status_text.set("Timelapse running")

    def _tl_stop(self):
        self._timelapse.stop()
        self._tl_start_btn.configure(state="normal", bg=C["orange"])
        self._tl_stop_btn.configure(state="disabled", bg=C["dgray"])
        count = self._timelapse.frame_count()
        self._tl_progress.configure(
            text=f"Stopped.  {count} frames captured.", fg=C["lgray"])
        self._status_text.set("Timelapse stopped")
        self._update_light_dot()

    def _tl_on_frame(self, idx: int, filepath: str):
        total     = self._tl_total.get()
        total_str = "∞" if total == 0 else str(total)
        def _update():
            self._tl_progress.configure(
                text=f"Running...  {idx} / {total_str} frames", fg=C["green"])
            self._tl_last_file.configure(text=f"Last: {os.path.basename(filepath)}")
        self.after(0, _update)

    def _tl_on_finish(self, total: int):
        def _update():
            self._tl_start_btn.configure(state="normal", bg=C["orange"])
            self._tl_stop_btn.configure(state="disabled", bg=C["dgray"])
            self._tl_progress.configure(
                text=f"Complete.  {total} frames captured.", fg=C["lgray"])
            self._status_text.set("Timelapse complete")
            self._set_light(self._pre_tl_brightness)
        self.after(0, _update)

    def _tl_on_error(self, msg: str):
        self.after(0, lambda: self._tl_progress.configure(
            text=f"Error: {msg}", fg=C["red"]))

    def _tl_on_light_on(self):
        pct = self._light.get_brightness()
        def _update():
            self._light_pct.set(pct)
            self._light_pct_lbl.configure(text=f"{pct:.0f}%")
            self._update_light_dot()
        self.after(0, _update)

    def _tl_on_light_off(self):
        def _update():
            self._light_pct.set(0.0)
            self._light_pct_lbl.configure(text="0%")
            self._update_light_dot()
        self.after(0, _update)

    # ══════════════════════════════════════════════════════════════════════════
    # Camera settings actions
    # ══════════════════════════════════════════════════════════════════════════

    def _on_ae_toggle(self):
        self._exp_entry.configure(
            state="disabled" if self._ae_var.get() else "normal")

    def _reset_cam_settings(self):
        """Reset to full-auto: AE on, AWB on, clear manual exposure/gain."""
        self._ae_var.set(True)
        self._awb_var.set(True)
        self._exp_var.set(str(DEFAULT_EXPOSURE_US))
        self._gain_var.set(DEFAULT_GAIN)
        self._on_ae_toggle()
        self._ae_btn.configure(text="ON", bg=C["green"])
        self._camera.apply_controls({"AeEnable": True, "AwbEnable": True})
        self._settings_status.configure(text="Reset to auto ✓", fg=C["green"])
        self._status_text.set("Camera reset to auto")
        self.after(3000, lambda: self._settings_status.configure(text=""))

    def _load_fixedlight_preset(self):
        """Load the exact camera settings from the original fixedlight.py."""
        self._ae_var.set(False)
        self._awb_var.set(False)
        self._exp_var.set(str(FIXEDLIGHT_PRESET["ExposureTime"]))
        self._gain_var.set(FIXEDLIGHT_PRESET["AnalogueGain"])
        # Reflect AE state in the entry widget
        self._on_ae_toggle()
        # Refresh the ON/OFF button labels on both toggles
        for var, btn in [(self._ae_var, self._ae_btn)]:
            btn.configure(text="ON" if var.get() else "OFF",
                          bg=C["green"] if var.get() else C["dgray"])
        # Apply immediately
        self._camera.apply_controls(FIXEDLIGHT_PRESET)
        self._settings_status.configure(
            text="Preset loaded and applied ✓", fg=C["green"])
        self._status_text.set("Preset applied")
        self.after(3000, lambda: self._settings_status.configure(text=""))

    def _apply_cam_settings(self):
        ctrl = {
            "AeEnable":     self._ae_var.get(),
            "AwbEnable":    self._awb_var.get(),
            "AnalogueGain": round(self._gain_var.get(), 2),
        }
        if not self._ae_var.get():
            try:
                exp = int(self._exp_var.get())
                if exp < 100 or exp > 10_000_000:
                    raise ValueError
                ctrl["ExposureTime"] = exp
            except ValueError:
                messagebox.showerror(
                    "Invalid exposure",
                    "Enter an exposure time between 100 and 10 000 000 µs.")
                return

        self._camera.apply_controls(ctrl)
        self._settings_status.configure(text="Settings applied ✓", fg=C["green"])
        self._status_text.set("Settings applied")
        self.after(3000, lambda: self._settings_status.configure(text=""))

    def _apply_pwm_freq(self):
        try:
            hz = int(self._pwm_freq_var.get())
            if hz < 10 or hz > 100_000:
                raise ValueError
        except ValueError:
            self._pwm_freq_status.configure(
                text="Enter a frequency between 10 and 100 000 Hz", fg=C["red"])
            return
        ok = self._light.set_frequency(hz)
        if ok:
            self._pwm_freq_status.configure(
                text=f"PWM frequency set to {hz} Hz ✓", fg=C["green"])
            self._status_text.set(f"PWM freq: {hz} Hz")
        else:
            self._pwm_freq_status.configure(
                text="Failed – see log for details", fg=C["red"])
        self.after(4000, lambda: self._pwm_freq_status.configure(text=""))

    def _reinit_light(self):
        """Tear down and re-initialise the LightController."""
        self._light.cleanup()
        self._light = LightController(prefer_sysfs=self._pwm_hw_var.get())
        self._timelapse.light = self._light
        self._on_light_slider(str(self._light_pct.get()))
        self._status_text.set(f"Light re-init: {self._light.backend}")
        log.info("Light re-initialised: %s", self._light.backend)

    # ══════════════════════════════════════════════════════════════════════════
    # Cleanup
    # ══════════════════════════════════════════════════════════════════════════

    def _on_close(self):
        log.info("Shutting down...")
        if self._timelapse.is_running():
            self._timelapse.stop()
        self._preview_stop.set()
        self._camera.close()
        self._light.turn_off()
        self._light.cleanup()
        self.destroy()


# ══════════════════════════════════════════════════════════════════════════════
# Entry point
# ══════════════════════════════════════════════════════════════════════════════

def main():
    app = dishcamApp()
    app.mainloop()


if __name__ == "__main__":
    main()
```
